# Realistic vOlumetric-Approach to Simulate Transcranial Electric Stimulation – ROAST – a fully automated open-source pipeline

**DOI:** 10.1101/217331

**Authors:** Yu Huang, Abhishek Datta, Marom Bikson, Lucas C. Parra

## Abstract

**Objective:** Research in the area of transcranial electrical stimulation (TES) often relies on computational models of current flow in the brain. Models are built based on magnetic resonance images (MRI) of the human head to capture detailed individual anatomy. To simulate current flow on an individual, the subject’s MRI is segmented, virtual electrodes are placed on this anatomical model, the volume is tessellated into a mesh, and a finite element model (FEM) is solved numerically to estimate the current flow. Various software tools are available for each of these steps, as well as processing pipelines that connect these tools for automated or semi-automated processing. The goal of the present tool – ROAST – is to provide an end-to-end pipeline that can automatically process individual heads with realistic volumetric anatomy leveraging open-source software and custom scripts to improve segmentation and execute electrode placement.

**Approach:** ROAST combines the segmentation algorithm of SPM8, a Matlab script for touch-up and automatic electrode placement, the finite element mesher iso2mesh and the solver getDP. We compared its performance with commercial FEM software, and SimNIBS, a well-established open-source modeling pipeline.

**Main Results:** The electric fields estimated with ROAST differ little from the results obtained with commercial meshing and FEM solving software. We also do not find large differences between the various automated segmentation methods used by ROAST and SimNIBS. We do find bigger differences when volumetric segmentation are converted into surfaces in SimNIBS. However, evaluation on intracranial recordings from human subjects suggests that ROAST and SimNIBS are not significantly different in predicting field distribution, provided that users have detailed knowledge of SimNIBS.

**Significance:** We hope that the detailed comparisons presented here of various choices in this modeling pipeline can provide guidance for future tool development. We released ROAST as an open-source, easy-to-install and fully-automated pipeline for individualized TES modeling.

## 1. Introduction

Models of current flow in the brain are important in research related to transcranial electrical stimulation (TES) as well as electroencephalography (EEG). TES modalities include trancranial direct current and alternating current stimulation (tDCS and tACS), which are generally limited to weak currents of no more than 2 mA. But TES also includes electroconvulsive therapy (ECT), which can go up to 800 mA (Guleyupoglu et al., 2013). In TES currents are applied to the scalp and modeling aims to determine which brain areas are stimulated (Datta et al., 2009; Lee et al., 2012), or where one should place electrodes to “target” a specific brain area (Dmochowski et al., 2011). In the case of EEG, currents generated by the brain are measured as voltage fluctuations on the scalp, and the objective of modeling is to determine the spatial origin of these brain currents. For this “source localization” in EEG (Haufe et al., 2011), one requires the identical current-flow models as for targeting TES (Dmochowski et al., 2017).

A multitude of current-flow models have been developed over the years with increasing level of detail (Rush and Driscoll, 1969; Ferdjallah et al., 1996; Stecker, 2005; Miranda et al., 2006; Datta et al., 2008; Dmochowski et al., 2012; Wagner et al., 2004). Starting in 2009 these models have been built based on individual head anatomy as captured with magnetic resonance imaging (MRI), e.g. with a T1-weighted image (Datta et al., 2009; Sadleir et al., 2010; Parazzini et al., 2011; Datta et al., 2012; Minhas et al., 2012; Datta et al., 2010; Wagner et al., 2007; Datta et al., 2011; Opitz et al., 2015). Today, the major steps for this modeling process include segmenting the MRI into different tissue compartments, assigning conductivity to each compartment, placing virtual electrodes on the model, tessellating this volumetric anatomy into a 3D mesh, and numerically solving the Laplace equation for the voltage distribution on this finite element model (FEM) (Figure 1).

**Figure 1:**
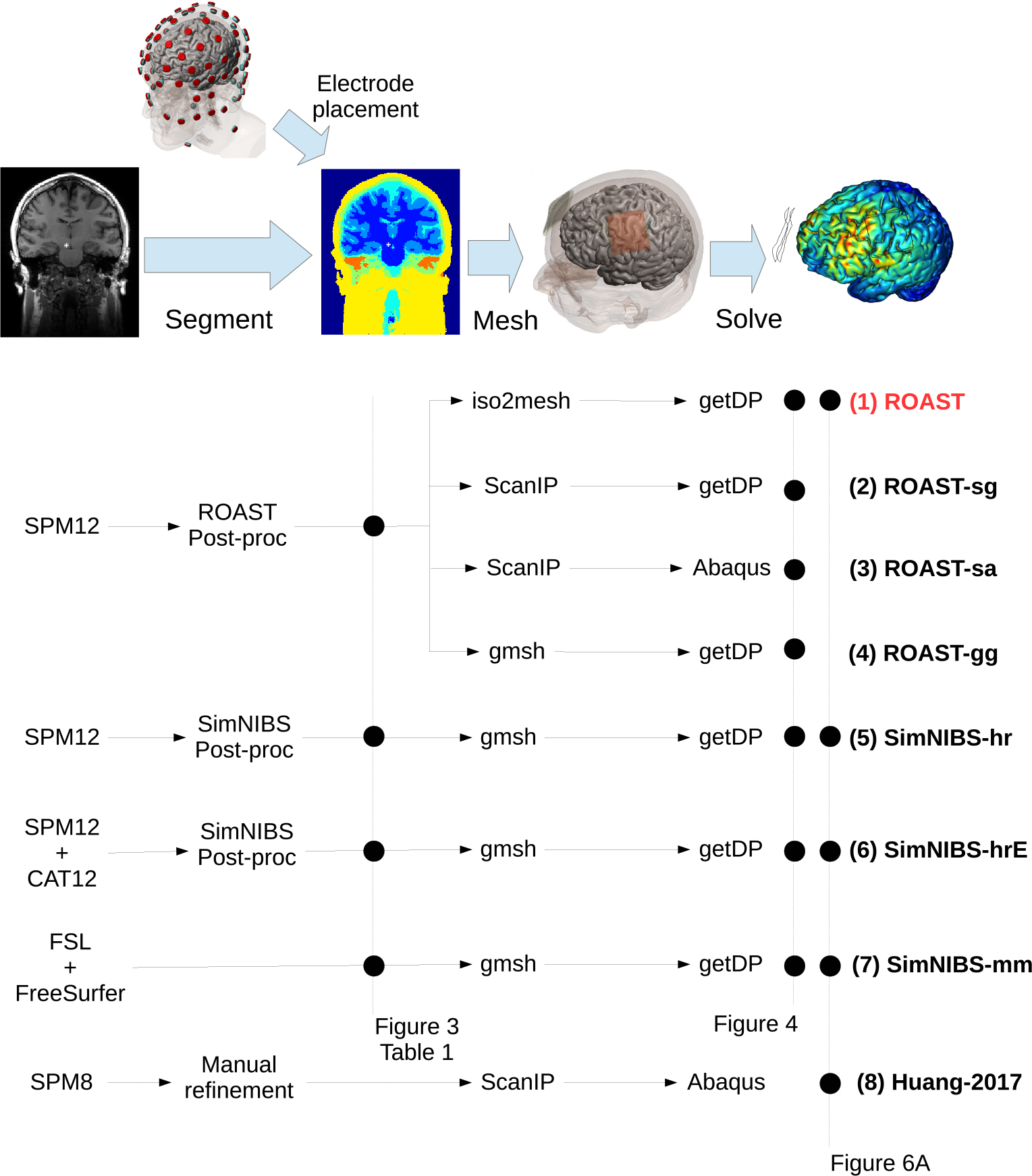
Candidate pipelines for building a current-flow model of the head. The input is the MRI of an individual, and the output of each pipeline is the predicted electric field distribution. The different pipelines we are evaluating are ROAST, with its variants (ROAST-sg, ROAST-sa, ROAST-gg), three different segmentation options in SimNIBS (SimNIBS-hr, SimNIBS-hrE and SimNIBS-mm), and the approach published in Huang et al. (2017). Data from different methods at different stages in these pipelines are used for comparison, as indicated by the black dots on the dashed vertical lines: Figure 3 / Table 1 – comparing segmentation; Figure 4 – comparing electric field distribution; Figure 6A – validating each pipeline using actual recordings.

**Table 1:** Segmented head tissues from each modeling software that are compared. Refer to Figure 1 for details on each pipeline. WM: white matter; GM: gray matter; CSF: cerebrospinal fluid.

| Pipelines | Segmented tissues |
| --- | --- |
| (1) ROAST | WM, GM, CSF, skull, scalp, air cavities |
| (5) SimNIBS-hr | WM, GM, CSF, eyes, skull, scalp, air cavities |
| (6) SimNIBS-hrE | WM, GM, CSF, eyes, ventricles, skull, scalp, air cavities |
| (7) SimNIBS-mm | WM, GM, cerebellum, CSF, ventricles, skull, scalp |

Various software tools are available to execute each of these steps. For example, SPM (Statistical Parametric Mapping, Friston (2007)), FSL (FMRIB Software Library, Smith (2002); Smith et al. (2004)) and FreeSurfer (Dale et al., 1999; Fischl et al., 1999) can all generate segmentations for the brain and head, with each one having different pros and cons (for a review on these segmentation tools, see e.g. Huang and Parra (2015)). A finite element mesh can be generated using open-source tools (e.g. iso2mesh (Fang and Boas, 2009), Gmsh (Geuzaine and Remacle, 2009)) or commercial software (e.g. ScanIP (Simpleware Ltd, Exeter, UK), Mimics (Materialise NV, Leuven, Belgium)). The same is true for FEM solvers (open-source: getFEM++ (Renard and Pommier, 2010), getDP (Dular et al., 1998); commercial: Abaqus (SIMULIA, Providence, RI), COMSOL Multiphysics (COMSOL Inc., Burlington, MA)). SciRun is another open-source tool that can generate meshes and solve the FEM (Weinstein et al., 1998; Fuchs et al., 1998). Given this great variety and complexity, there is a need to automate these tools into a complete and easy-to-use processing pipeline. Existing pipelines are either not fully automated (e.g. NFT (Acar and Makeig, 2010)) or difficult to use (e.g. SciRun (Dannhauer et al., 2012)). Notable in terms of ease of use (if not of installation) is SimNIBS (Windhoff et al., 2011; Thielscher et al., 2015; Saturnino et al., 2015), which integrates FreeSurfer, FSL, Gmsh and getDP to provide a complete end-to-end solution. In the TES modeling literature, volumetric finite element models are preferred over surface-based boundary element methods (BEM) that are more common for EEG modeling (e.g. Fuchs et al., 1998). While a BEM can capture the detailed gyral/sulcal surfaces, it has limited abilities to represent the anatomical morphology. For instance, boundary surfaces between tissues need to be entirely contained within one another, which makes it difficult to implement anatomical details, such as the optic foramen, which is a highly conducting conduit into the skull. Moreover, generating surface segmentations for BEM (Dale et al., 1999) is computationally demanding, compared to volumetric segmentation algorithm based on probability inferences (Ashburner and Friston, 2005). Recently SimNIBS V2.1 was released (Nielsen et al., 2018) which incorporates the volumetric segmentation approach of SPM. This greatly reduces the computation time for segmentation from ~ 10 hours to 40 minutes. However, during post-processing of SPM-generated segmentation, SimNIBS still converts volumetric segmentation into surfaces, which limits the flexibility of volumetric modeling of anatomic details such as the optic foramen (see Section 3.1 for details).

Here we present a fast automated pipeline that operates entirely with volumes which we released in 2017 under the code name ROAST, a short hand for “Realistic vOlumetric-Approach to Simulate Transcranial Electric Stimulation”. In its current release ROAST uses the segmentation algorithm of SPM version 12 and applies it to the entire head and neck (Huang et al., 2013). We integrate this with our post-processing routine that ensures continuity of the cerebrospinal fluid (CSF) and skull and an additional tool that automatically places virtual electrodes on the volumetric segmentation. This is then followed by meshing with iso2mesh and FEM solving with getDP (Figure 1). Finally, we use volumetric visualization of the resulting electric fields. ROAST is based on Matlab but is otherwise entirely open source. It processes individual MRI volumes in a fully automated fashion to generate 3D renderings of the resulting electric field distributions. The user can specify any number of electrodes within the 10-05 and/or BioSemi-256 system. The end-to-end processing time is typically less than 30 minutes, which is much faster than alternative approaches (e.g. SimNIBS). It is also significantly easier to use and to install as compared to other tools (e.g. SCIRun).

Initial evaluation of ROAST compared to commercial software (ScanIP and Abaqus) and SimNIBS V2.0 on the MNI-152 standard head (Grabner et al., 2006) was published before (Huang et al., 2018). Here we extend the analysis by also comparing the segmentation results of ROAST and SimNIBS to a hand-labeled segmentation of an individual head which serves as ground truth. We also add the newer version of SimNIBS (V2.1) to the comparison of the electric fields published in Huang et al. (2018). We further extend the comparison between ROAST and SimNIBS in terms of segmentation and simulated electric field to 14 human heads published previously (Huang et al., 2016). To our knowledge this is the first quantitative comparison of these TES modeling tools. This evaluation reveals some difference in the predicted field distribution between ROAST and various versions of SimNIBS. To determine which of these is closer to actual field measures in human heads we also compare the predictions to the recorded data (Huang et al., 2016). ROAST and SimNIBS do not seem to differ from each other in prediting the electric field distribution, but one has to know sufficient details about SimNIBS options to avoid it failing on some of the subject heads. The latest release of ROAST is freely available and we hope that this can make current-flow models accessible to a broader group of researchers and clinical investigators.

## 2. Methods

### 2.1. Components of ROAST pipeline

ROAST stands for Realistic vOlumetric-Approach to Simulate Transcranial Electric Stimulation. It is composed of the following components:

- Segmentation The “Unified Segmentation” algorithm (Ashburner and Friston, 2005) implemented in SPM12 is used, with an extended tissue probability map covering the entire head (Huang et al., 2013). This allows one to model the head with a field-of-view (FOV) that exends down to the neck. Segmentation from SPM12 is further improved by a Matlab script. This post-processing attempts to smooth the segmentation, fill holes on the cerebrospinal fluid (CSF), and remove the disconnected voxels. The first part of this post-processing has been presented previously (Huang et al., 2013), and it is based on morphological operations. The parameters in these operations are selected conservatively and thus gaps in the thin layers of gray matter, CSF and skull cannot be fully removed. Here, these remaining gaps were filled using simple heuristics (e.g., to fill gaps in CSF, we check if any brain voxel touches skull/scalp/air, if so, then we convert the skull/scalp/air neighbors to CSF). While this fills in all gaps in gray matter and CSF, we do not apply this process indiscriminately to all of the skull, as we want to preserve skull openings in the optic canal and the foramen magnum which can serve as important conduits of currents into the brain. To preserve these genuine openings in the skull, a special mask was made marking these regions in the tissue probability maps used by the SPM12 segmentation routine and this mask was mapped into the individual MRI space during segmentation. These regions were then treated as exceptions during the patching process.
- Electrode placement This is a Matlab script we developed based on previous work (Huang et al., 2013). In its latest version, users can place virtual electrodes freely among the locations of the 10-20, 10-10, 10-05 or BioSemi-256 EEG system. They can also place the electrode at any arbitrary location on the scalp by providing the coordinates in the MRI voxel space. Furthermore, the shape of each electrode can be selected to be either a disc, pad or ring, with customizable size and orientation (for pad).
- Finite element meshing The Matlab toolbox iso2mesh (Fang and Boas, 2009) is used to generate the volumetric mesh. Specifically, the “cgalv2m” function was used to generate a tetrahedral mesh directly from the segmented MRI. This is made possible by the CGAL package (Rineau et al., 2009), which is capable of generating a volumetric mesh from 3D multi-domain images. Users can customize the mesh options provided by iso2mesh, such as the element size.
- FEM solving The open-source solver getDP (Dular et al., 1998) is used to solve the underlying Laplacian equation (Griffiths, 1999). Users specify how much current goes into each anode and out of each cathode (Neumann boundary condition). The tissue conductivity can be customized, with the default based on previous literature (Crille et al., 1922; Burger and van Milaan, 1943; Freygang Jr and Landau, 1955; Ranck Jr., 1963; Hasted, 1973; Geddes, 1987; De Mercato and Garcia Sanchez, 1992; Gabriel, 1996; Akhtari et al., 2002).

Figure 1 shows the ROAST pipeline (text highlighed in red). ROAST is currently available online at https://www.parralab.org/roast/. A README file there contains more details on the features of this software. As a fully-automated and easy-to-use pipeline, users do not have to install separate packages. They only need to install Matlab, download ROAST, and enter a one-line command that selects the desired MRI (in NIfTI format) and the desired electrode locations with the amount of injected current. A simulation result will then be generated within 20–30 minutes for a typical 1 mm resolution head MRI (tested on a typical dual-core computer with 16 GB memory).

### 2.2. Comparison of different modeling pipelines

We evaluated the free ROAST pipeline along with variants that use commercial software (Huang et al., 2017), as well as the free SimNIBS pipeline (Windhoff et al., 2011; Nielsen et al., 2018), and various intermediate version (Figure 1). This leads to a comparison across four different segmentation methods (Figure 3, Table 1) and seven different TES modeling workflows (Figure 4).

**Figure 2:**
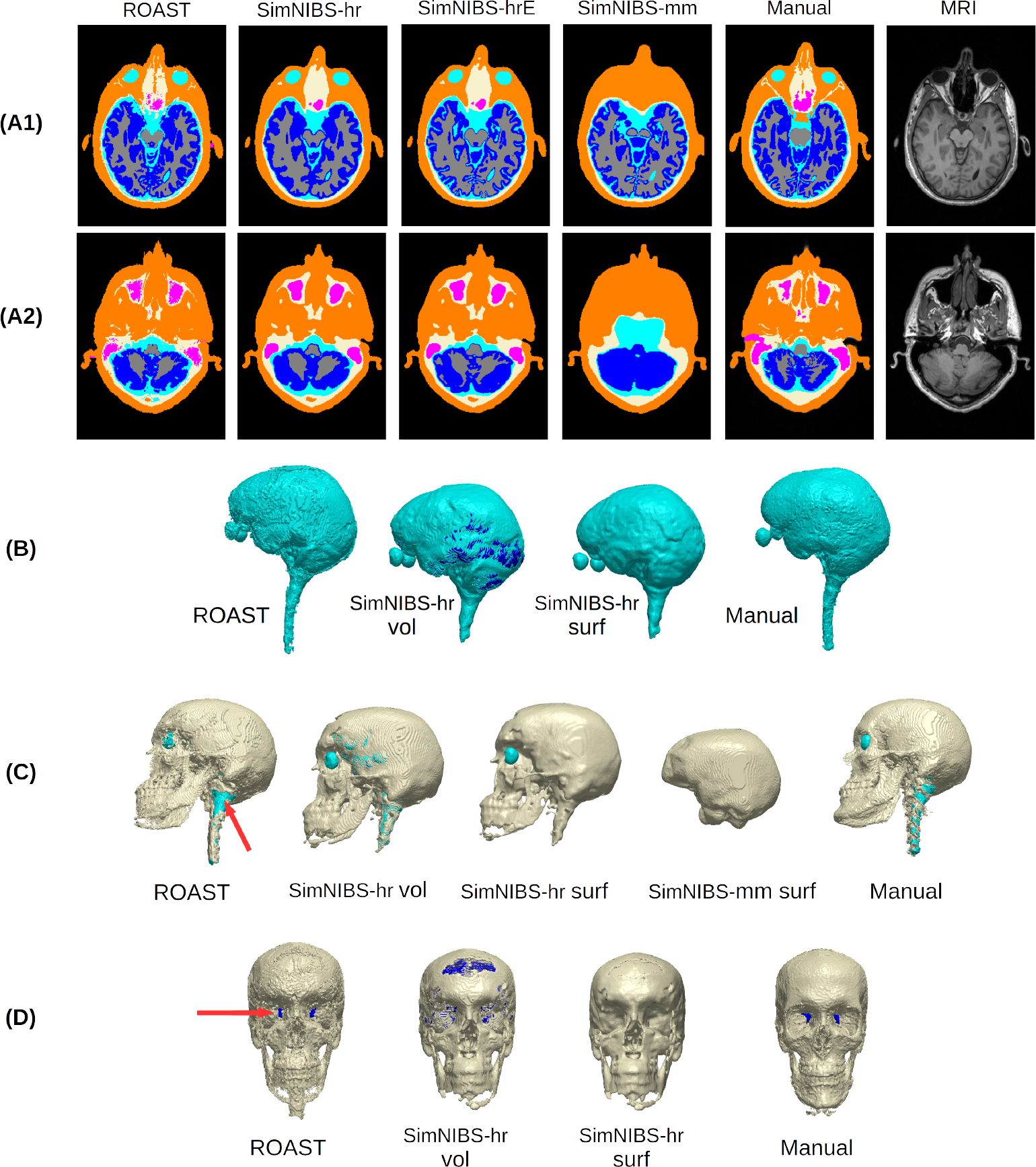
Example brain slices (A1, A2) and 3D renderings (B–D) showing the segmentation from different software tools (ROAST, SimNIBS-hr, SimNIBS-hrE, SimNIBS-mm) and the manual segmentation (Manual) of the individual head S1. For the 3D renderings, both the volumetric (vol) and surface (surf) formats are displayed for the CSF and skull generated by SimNIBS-hr. Red arrows indicate the optic foramen and the foramen magnum. Note that SimNIBS-mm is missing portions of midbrain and spinal cord due to is limited FOV. Refer to Figure 1 for the details of the four segmentation methods.

**Figure 3:**
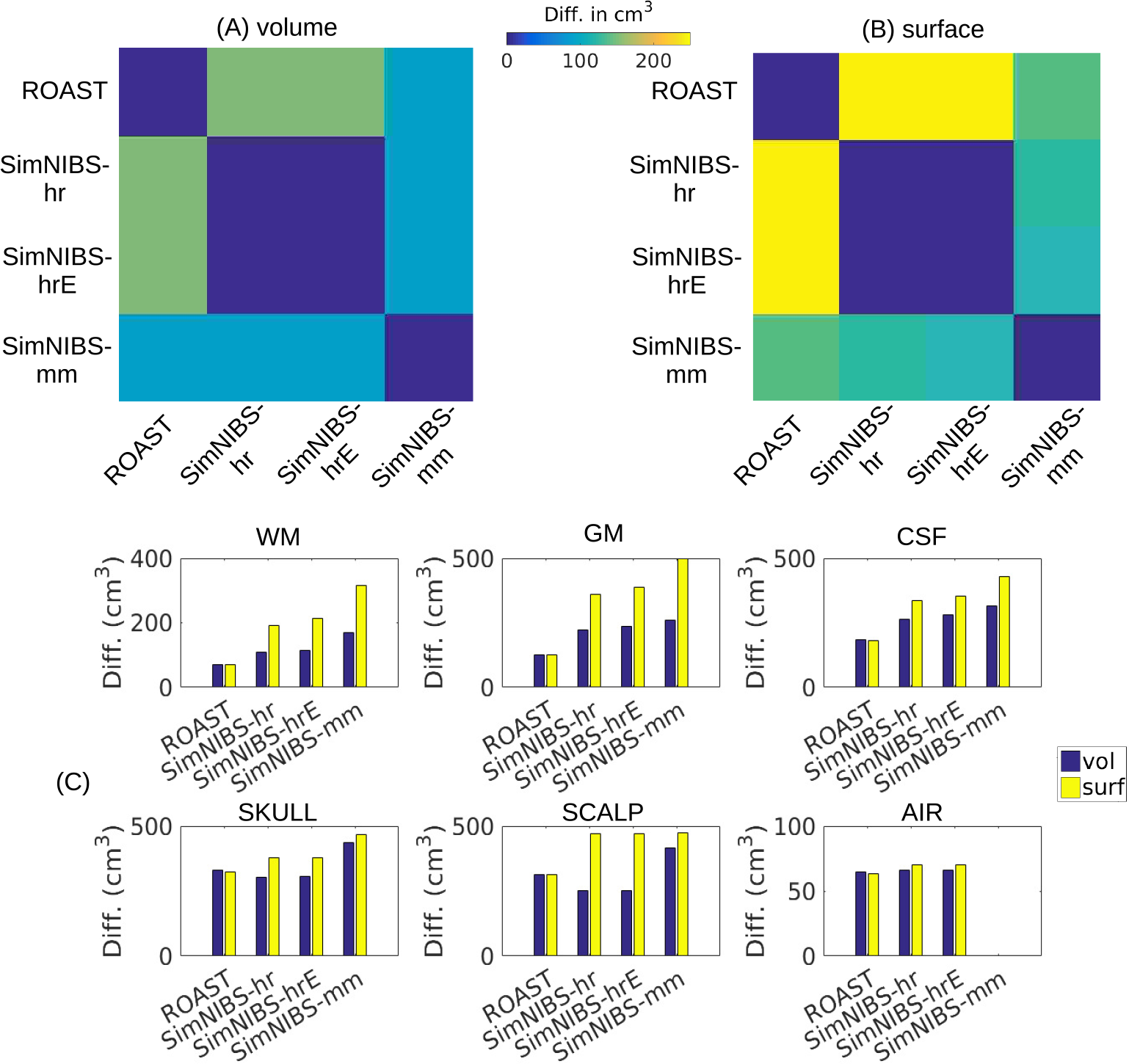
The absolute volumes (cm^3^) of the segmentation differences between the four methods on the overall segmentation of the individual head (S1) are displayed as matrices for volumetric (A) and surface (B) format. Also the difference for each tissue type for each method compared against the manual segmentation is shown as bar plots (C), also for the volumetric and surface formats. The “surf” format (yellow bars) for ROAST is in fact the upsampled segmentation results from ROAST. Refer to Figure 1 and Section 2.2.1 for the details of the four segmentation methods. WM: white matter; GM: gray matter; CSF: cerebrospinal fluid.

**Figure 4:**
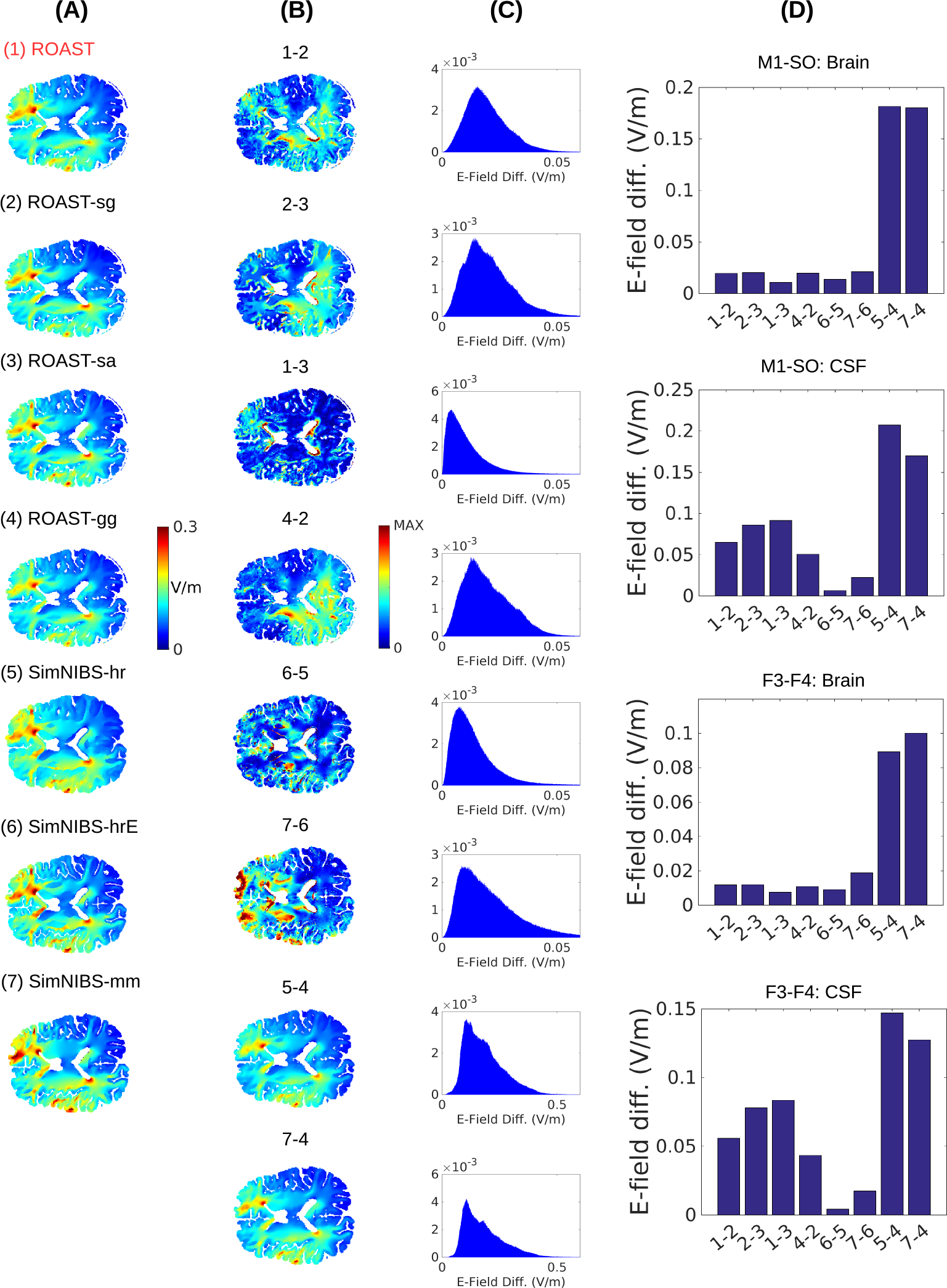
(A) Magnitude of electric field under stimulation montage M1-SO predicted by various modeling pipelines for the individual head S1. Refer to Figure 1 for the details on the different modeling pipelines. The voxelwise differences between these methods as calculated by Eq. 1 are shown for the same 2D slice (B) and as histograms (C) for the entire brain volume including cerebellum. The “MAX” value in the color axis of (B) can be found in the corresponding histogram in (C). The average differences in the brain and CSF for montage M1-SO and F3-F4 are shown as bar plots in (D). The numbers in the text below each bar correspond to the numbers given to each method above each slice view. Refer to the main text for the explanations on each pair of comparison between these methods.

#### 2.2.1. Segmentation

A T1-weighted MRI of an adult male subject (S1) published before (Huang et al., 2013) was used to evaluate the performance of different software on segmentation. The reason for using this individual is that it has been manually segmented from scratch (Datta et al., 2009) so there is no bias introduced by initializing the manual segmentation with the results from an automated segmentation software.

The first segmentation candidate was obtained by ROAST (see Section 2.1 for details). ROAST saves the segmentation results in the voxel space of the MRI. The other three candidates for comparison are part of the SimNIBS toolbox (headreco, headrecoE, mri2mesh; here for simplicity we abbreviated them as SimNIBS-hr, SimNIBS-hrE and SimNIBS-mm, respectively). The earlier segmentation routine in SimNIBS is SimNIBS-mm, which combines FreeSurfer (Dale et al., 1999; Fischl et al., 1999) and FSL (Smith, 2002) to segment the brain and non-brain tissues, respectively (Windhoff et al., 2011). Note that this routine cannot segment any head MRI with size bigger than 256 pixels in any direction. As S1 has a size of 280×320×208, we cut it into 246×256×208 by removing the empty slices in the posterior direction and also the slices of the lower part of the head. Also, SimNIBS-mm saves segmentation results in the FreeSurfer standard space, so we registered and resampled that back into the MRI voxel space to be consistent with the other approaches.

Recently a newer version of SimNIBS was released that incorporates SPM12 as its segmentation function (Nielsen et al., 2018), and uses the extended atlas published in Huang et al. (2013) as the prior probability map for segmentation (this map covers the lower part of the head). With this, SimNIBS can now segment heads with sizes bigger than 256 pixels in any dimension, and runs in 1 to 2 hours instead of 10 hours. The entire modeling process was named headreco, and we name it as SimNIBS-hr. SimNIBS-hr also provides an option to use the CAT12 (Computational Anatomy Toolbox 12, http://www.neuro.uni-jena.de/cat/) to enhance cortical segmentation by capturing sulci in more detail. We denote the segmentation from this as SimNIBS-hrE (E meaning enhanced).

In SimNIBS-mm, the head segmentation generated by FreeSurfer and FSL is in surface format and is used directly for the subsequent mesh generation (Figure 1, Pipeline 7); in SimNIBS-hr and SimNIBS-hrE, SPM12 is used to generate volumetric segmentation which is processed by SimNIBS and converted into surface format for the mesh generation (Figure 1, Pipelines 5 and 6). For all these routines, SimNIBS saves the surface segmentation (.stl) right before entering the mesher, and it also saves this same segmentation data in volumetric format (.nii) by just voxelizing the surfaces. These segmentation data are saved under the `m2m_*` folder. ROAST outputs segmentation in a volumetric format (.nii). The detailed tissue segmentations for each of these four software are summarized in Table 1.

To find out how volumetric and surface formats affect the segemntation results, we compared both SimNIBS formats to the segmentation generated from ROAST, as well as the manual segmentation as the ground truth. For the volumetric segmentation from SimNIBS, we performed the following operations to make them consistent with head segmentation from ROAST (see Table 1): for SimNIBS-hr, combine eyes with CSF as CSF segmentation from ROAST includes eyes; for SimNIBS-hrE, combine eyes and ventricles with CSF, as ventricles here are output as a separate mask; for SimNIBS-mm, combine ventricles with CSF, and combine cerebellum with gray matter, as in the other three routines gray matter always includes cerebellum; note air cavities are labeled as skull by SimNIBS-mm. For the surface format of SimNIBS-genarated segmentation, we converted the surfaces to volumes and upsampled the volumes from 1 mm to 0.2 mm resolution to ensure that surfaces are still fully closed in voxelized grid (ROAST and manual segmentation were also upsampled to 0.2 mm for comparison). This was done in ScanIP and the same operations as described above to combine tissues were performed on the upsampled volumes from SimNIBS. Note that for SimNIBS-mm we registered the converted volumes back into MRI voxel space before the upsampling.

As the differences of these segmentation mainly occur at the tissue boundaries (Figure 2), a conventional similarity measure such as the Dice coefficient (Dice, 1945) will not meaningfully capture the differences. Therefore, here we report the absolute volumes (in the unit of cm^3^) of the segmentation differences between different methods and against the ground truth (i.e., hand-segmented tissue masks). This was done on both the volumetric and surface format of the segmentation (for ROAST and manual segmentation, the surface format is simply the upsampled version of volumetric data). When computing the difference between SimNIBS-mm and other methods we excluded voxels that are outside the FOV of mri2mesh routine.

To evaluate the differences between the four segmentation routines in Table 1 on a larger dataset, we performed the same analysis on the 14 subjects previously published (Huang et al., 2016). As the manual segmentation of these heads were initialized using SPM software, they are biased and thus cannot be used as the ground truth. Therefore we instead compared across different pairs of the four methods and only focused on the volumetric format.

#### 2.2.2. Simulation of electric field

On the same subject (S1), we compared the seven different workflows in Figure 1 in terms of the predictions on electric field distribution. Two different electrode montages were used for all the pipelines: M1-SO, a typical configuration used in most TES studies to stimulate the motor cortex (Datta et al., 2012); and F3-F4, a montage frequently used in studies of cognitive function (Seibt et al., 2015). In ROAST and SimNIBS electrodes were all modeled as small discs with radius and height being 6 mm and 2 mm, respectively. Gel of 2 mm thickness was also added under each electrode. The whole surface of the electrode was treated as the connector.

To test how different FEM software perform on the same segmentation data, we run the models using different combinations of either open-source or commercial FEM meshers and solvers, but identical segmentation and automated electrode placement routine (ROAST, ROAST-sg, ROAST-sa, ROAST-gg). For ROAST, iso2mesh and getDP serve as the mesher and solver, respectively (see Section 2.1 for details). In ROAST-sg, adaptive meshing (ScanFE-Free algorithm) was used in ScanIP and the output mesh was converted to .msh format that is compatible with getDP. ROAST-sa essentially follows the same details as described in Huang et al. (2013), which uses commercial software ScanIP and Abaqus. To compare how SimNIBS-generated segmentation affects the modeling results compared to SPM12-generated segmentation, we feed the segmented masks from SPM12 into gmsh and getDP, leading to ROAST-gg. Since gmsh only accepts surface segmentation as its input format (Geuzaine and Remacle, 2009), the volumetric masks from SPM12 were first converted into .stl format using iso2mesh in ROAST-gg before entering gmsh. For these four pipelines, the same conductivity values are used as in Huang et al. (2013), and the boundary condition was set as 1 mA current injection at the anode (note this 1 mA is calculated precisely by using the exact anode area calculated from the tetrahedral mesh elements).

The 5th to 7th pipelines in Figure 1 are three different segmentation options in SimNIBS (Version 2.1): headreco (SimNIBS-hr), headreco with CAT12 toolbox (denoted SimNIBS-hrE here) and mri2mesh (SimNIBS-mm) (Windhoff et al., 2011; Nielsen et al., 2018). For these three pipelines, electrode placement was done in SimNIBS graphic user interface (GUI) by selecting the names of the corresponding electrodes. Same tissue conductivity values were entered in the GUI, and the injected current was also set to 1 mA at the anode in the GUI. The model was then generated and solved using the built-in tools gmsh and getDP, respectively (Windhoff et al., 2011).

For the ease of comparing the outputs from these pipelines, the solutions on the mesh grid were resampled onto a regular grid with the same dimensions and resolution as the original MRI. For the ROAST results (1st to 4th pipelines in Figure 1) this was done using Matlab function `TriScatteredInterp()`. For the SimNIBS results (5th to 7th pipelines in Figure 1) this was done automatically by turning on the “Interpolate to a nifti volume” option in SimNIBS, and then registering and reslicing back to the original MRI voxel space. Note that SimNIBS-mm saves the results in the FreeSurfer standard space. These results were thus registered and resliced into the original voxel space of the input MRI. The voxelwise absolute differences of the electric field distribution were calculated between two methods A and B:

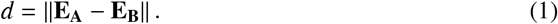

Here **E** indicates the electric field. We compare the results of the open-source tools to the commercial FEM software and the results using the SPM12 segmentation to the SimNIBS segmentation. Segmentation masks from each modeling pipeline were used to extract the tissue-specific electric field values generated by that pipeline. When comparing electric fields between two methods (e.g., ROAST-gg vs. SimNIBS-mm, see Figure 1), only locations with the same segmentation labels were considered.

To evaluate the differences between ROAST and SimNIBS on a larger dataset, we also compared the predicted electric field on the 14 subjects previously published (Huang et al., 2016). Note here we excluded subject P06 as SimNIBS flipped the electric field along the left-right direction in this subject when interpolating the results onto the nifti volume.

### 2.3. Validation of modeling pipelines on in-vivo, intracranial measurements of electric potentials

When we compare across the seven pipelines in Section 2.2.2, there is no ground truth to evaluate which pipeline provides more accurate prediction on electric field distribution. To this end, we validated the major four automated pipelines against *in-vivo*, intracranial electrical recordings in human (Huang et al., 2016), and compared also to the manual method of Huang et al. (2017). This gives five sets of results (ROAST, SimNIBS-hr, SimNIBS-hrE and SimNIBS-mm, Huang-2017). Computational models were built for 14 subjects (P03, P04, P05, P06, P07, P08, P09, P010, P011, P013, P014, P015, P016, P017) using their pre-surgical MRIs, i.e. the MRI that does not capture the craniotomy or implanted electrodes. As the recorded data in Huang et al. (2016) are calibrated into 1 mA current injection, all the models solved in the four automated pipelines were set to 1 mA current injection as well. Stimulation electrodes were placed on the scalp according to photos of the subjects during the recording. All the models were run using the subject MRIs in ROAST and SimNIBS out-of-box, except for P04 and P010. For these two subjects specific options were used to make SimNIBS work properly. Specifically, option `“-v 1.5”` was used for P04 when running SimNIBS-hrE to increase the vertex density so that different tissue surfaces can be fully decoupled. Option `“-d no-conform”` was used for P010 when running SimNIBS-hr and SimNIBS-hrE to avoid complication when processing the MRI of P010 which has anisotropic resolution. Also for subject P014 we excluded one recording electrode which had an outlier value when solving with SimNIBS-hr and SimNIBS-hrE. For each subject, the model-predicted values and the recorded values were compared in terms of two criteria: (1) Pearson correlation coefficient *r* between recorded and predicted values; (2) the slope *s* of the best linear fit with predicted value as “independent” and measurement as “dependent” variables. The correlation captures how well the model predicts the distribution patterns of the electric fields, regardless of a potential mismatch in overall magnitude. The slope measures how accurate the model estimates the absolute magnitudes. Electric fields are calculated between adjacent recording electrodes, i.e. they represent the projection of the actual field vector in this direction – as the cosine it will be typically smaller than the absolute magnitude. The voxel coordinates of the recording electrodes were used to read out the predicted voltages from the models. See Huang et al. (2017) for details on the methodology. Again note that for SimNIBS-mm, those heads with more than 256 axial slices were cut into 256 axial slices (discard the lower portion of the head). Also, results from SimNIBS were registered and resliced into the original voxel space of the input MRI.

## 3. Results

The following three studies report results on segmentation, field modeling, and compare these against measured fields. Specifically, Section 3.1 analyzes segmentation results on a single high-quality manual segmentation using unbiased truth data. It focuses on comparing the options available for segmentation. Section 3.2 analyzes the resulting field distributions with different FEM options. Section 3.3 compares fields with empirical data from 14 *in-vivo*, intracranial recordings in human.

### 3.1. Segmentation performance

To evaluate segmentation against “ground truth” we use a head that has been segmented entirely by hand at 1 mm^3^ resolution and refined over the years (Datta et al., 2009; DaSilva et al., 2015; Mourdoukoutas et al., 2018). Segmentation in ROAST is based on SPM12, followed by custom post-processing to implement morphological constraints that are important for current-flow modeling, such as continuous CSF (see Section 2.1). SimNIBS offers three segmentation options based on SPM12, CAT12 and FSL/FreeSurfer (see Figure 1 and Section 2.2.1), which are followed by custom post-processing. In SimNIBS the post-segmentation processing enforce that one tissue type is fully enclosed by another. The result of these four different approaches are shown in Figure 2 for an individual head along with a segmentation performed entirely by hand (manual). Inspection of two axial slices (Figure 2A1–A2) suggests that the SimNIBS tools (SimNIBS-hr, SimNIBS-hrE, SimNIBS-mm) give smoother segmentations compared to ROAST, especially for the thin surfaces of CSF and skull. In contrast, ROAST gives more detailed segmentation on small structures such as the small bones around the eyes and nose, which is in agreement with the manual segmentation. The CAT12 toolbox appears to enhance the segmentation of the brain by giving more detailed gyri and sulci (comparing SimNIBS-hrE with SimNIBS-hr). SimNIBS-mm does not capture the eyeballs or any realistic skull structures around the eyes and nose (also see Panel C SimNIBS-mm surf). It segments the brain well with better gyrations compared to SimNIBS-hr and ROAST, but failed to separate the gray and white matter in the cerebellum.

We also show 3D renderings of the segmentation of the CSF and skull in Figure 2. For SimNIBS-hr, we show both the volumetric and the surface format, as indicated by SimNIBS-hr vol and SimNIBS-hr surf, respectively. This is because SimNIBS can save the segmentation in both volumetric and surface format (see Section 2.2.1 for details), but uses the surface segmentation for making the FEM. ROAST only operates in volumetric format. In the volumetric segmentation we often find gaps in thin layers of CSF and skull (as seen in the blue color shining through in Figure 2B–D). SimNIBS closes all these gaps when converting the volumetric segmentation from SPM12 into surfaces, but in the process it loses details of the real anatomy especially for the skull (compare SimNIBS-hr vol and SimNIBS-hr surf in Figure 2B–D). For example, the veridical openings of the skull such as the optic foramen and the foramen magnum (indicated by the red arrows in Figure 2CD) are all closed in the surface format. ROAST, on the other hand, meshes the volumetric segmentation directly. Consistent with the manual segmentation, the automated post-segmentation script in ROAST avoids filling in gaps at known foramen locations (Figure 2CD under ROAST). Overall the manual segmentation is smoother (e.g. for the CSF; Figure 2B) yet contains in some places more details than the automated methods (e.g. for facial bones and temple; Figure 2C). This specific balance between smoothness and details may require more sophisticated anatomic prior knowledge than what is currently used by the automated methods.

The absolute volumes in the unit of cm^3^ of the segmentation difference between all methods are shown as a matrix in Figure 3. We choose this measure instead of the Dice coefficients to quantify the segmentation differences because the segmentation mainly differs at tissue boundaries, which cannot be accurately captured by the Dice coefficients. We compute this difference measure before and after the volume-to-surface conversion of SimNIBS (Panel A and B respectively). There is, on this head, almost no difference in results between SimNIBS-hr and SimNIBS-hrE, but these two differ from the segmentations of ROAST and SimNIBS-mm. The SimNIBS post-segmentation routine that converts volumes into surfaces introduces additional discrepancies (compare Panel A and B in Figure 3). In addition, the different methods are compared against the manual segmentation (shown as bar plots for each specific tissue type in Figure 3C). The main observation here is that for all methods the largest discrepancies to manual segmentation are observed for gray matter, CSF and skull (all three compartment are thin layers and thus difficult to establish conclusively). ROAST has the best performance for white matter, gray matter and CSF. SimNIBS-mm has overall lower performance for all tissues compared to the other three methods. The volume-to-surface conversion causes a small drop in performance for all SimNINBS methods. However, we should note that factors such as continuity of a compartment may be ultimately more important than small changes in segmentation volumes. This may become more apparent when evaluating the resulting field distributions (see Section 3.2).

Segmentation evaluation on a larger dataset (Huang et al., 2016) is shown in Figure 6B. See Section 3.3 for details.

### 3.2. Electric fields predicted by different modeling pipelines

When computing the electric field distributions we wanted to evaluate the effect of different meshing and FEM solving toolboxes including free and commercial software. We do this for the segmentation results of ROAST. We also evaluated the results with the same free mesher and solver but for the different segmentation options of SimNIBS. This results in seven different pipeline options (see Figure 1).

The spatial distributions of electric field magnitudes in the brain under M1-SO montage are visually quite similar across these pipelines (Figure 4A). Note we ran the simulations on the same head as used in Section 3.1. We compared eight pairs of pipelines for the electric field in the brain and CSF:

- ROAST (1) vs ROAST-sg (2): shows the difference introduced from using open-source mesher iso2mesh instead of the commercial ScanIP;
- ROAST-sg (2) vs ROAST-sa (3): gives the difference between the free solver getDP and commercial solver Abaqus;
- ROAST (1) vs ROAST-sa (3): captures the difference between iso2mesh/getDP and ScanIP/Abaqus;
- ROAST-gg (4) vs ROAST-sg (2): gives the difference between gmsh and ScanIP;
- SimNIBS-hrE (6) vs SimNIBS-hr (5): shows the effect of adding the CAT12 toolbox with improved brain segmentation;
- SimNIBS-mm (7) vs SimNIBS-hrE (6): is the difference between SimNIBS version 2.0 and version 2.1;
- SimNIBS-hr (5) vs ROAST-gg (4): represents the difference from using SimNIBS-hr and ROAST segmentation, which shows the genuine difference between ROAST and SimNIBS (see below);
- SimNIBS-mm (7) vs ROAST-gg (4): shows the difference between the segmentation generated by SimNIBS-mm and ROAST.

For all these comparisons, the voxelwise difference of electric field as calculated by Eq. 1 for montage M1-SO is shown in the same brain slice in Figure 4B, with the histograms of the differences across the entire brain including the cerebellum shown in Figure 4C. The differences are generally smaller than 0.05 V/m except for the last two, which are between SimNIBS-hr / SimNIBS-mm and ROAST-gg. The average differences in the brain and CSF are shown for each pair of comparisons and each of the two stimulation montages in Figure 4D.

In terms of the electric field inside the brain it appears that using free vs. commercial mesher/solver does not significantly impact the result (Figure 4D, Brain, the first four comparisons). Within SimNIBS the different segmentation functions also give differences smaller than 0.03 V/m in the brain (Figure 4D, Brain, 6 vs. 5, 7 vs. 6). However, the differences shoot up to ∼0.1 V/m if one uses SimNIBS instead of ROAST (Figure 4D, Brain, 5 vs. 4, 7 vs. 4). The big difference between SimNIBS-mm (7) and ROAST-gg (4) can be explained by the different segmentation approaches used in the two software: ROAST is based on the SPM12 segmentation algorithm that works on voxelized image data (Ashburner and Friston, 2005). SimNIBS-mm, which utilizes FSL (Smith, 2002; Smith et al., 2004) and FreeSurfer (Dale et al., 1999; Fischl et al., 1999), generates a segmentation in the format of a surface mesh. However, this cannot explain the large difference between SimNIBS-hr (5) and ROAST-gg (4), as the two utilize exactly the same toolboxes for segmentation, meshing and solving (see Figure 1). The source of this difference lies in the post-processing of the segmentation. When building the mesh, SimNIBS converts the volumetric segmentation from SPM12 into decoupled surfaces (see Section 3.1, Figure 2 and Figure 3 for details). In practice, however, this big difference between ROAST-gg and SimNIBS-hr is not important as we do not distribute ROAST-gg. In fact, direct comparison between SimNIBS-hr and ROAST (5 vs. 1) does not show that much difference in the brain (Figure 6C).

The electric field inside the CSF is another interesting quantity to look at. Compared to the brain, the difference in the CSF is higher if getDP is used instead of Abaqus (Figure 4D, CSF, 2 vs. 3, 1 vs. 3). This is expected as the CSF is a very thin layer with a high jump of conductivity from its neighboring tissues (CSF: 1.65 S/m; gray matter: 0.276 S/m; skull: 0.01 S/m). Due to this jump in conductivity, computation of electric field from voltages across the tissue boundaries is analytically not defined and it depends on the numerical details of the solvers on how to compute its approximation from the resulting voltages (Engwer et al., 2017). This can be seen in Figure 5 where we take out a line in one example slice of the head and plot the electric field magnitude obtained from getDP and Abaqus. The field magnitudes are exactly the same inside the brain but are different on the boundaries of brain, CSF and skull (indicated by arrows in Panel B) despite nearly identical computed voltages in both solvers. Similarly to the comparison for the brain, the three different functions within SimNIBS also give differences smaller than 0.03 V/m (Figure 4D, CSF, 6 vs. 5, 7 vs. 6), but higher differences are found when comparing SimNIBS results with ROAST (Figure 4D, CSF, 5 vs. 4, 7 vs. 4).

**Figure 5:**
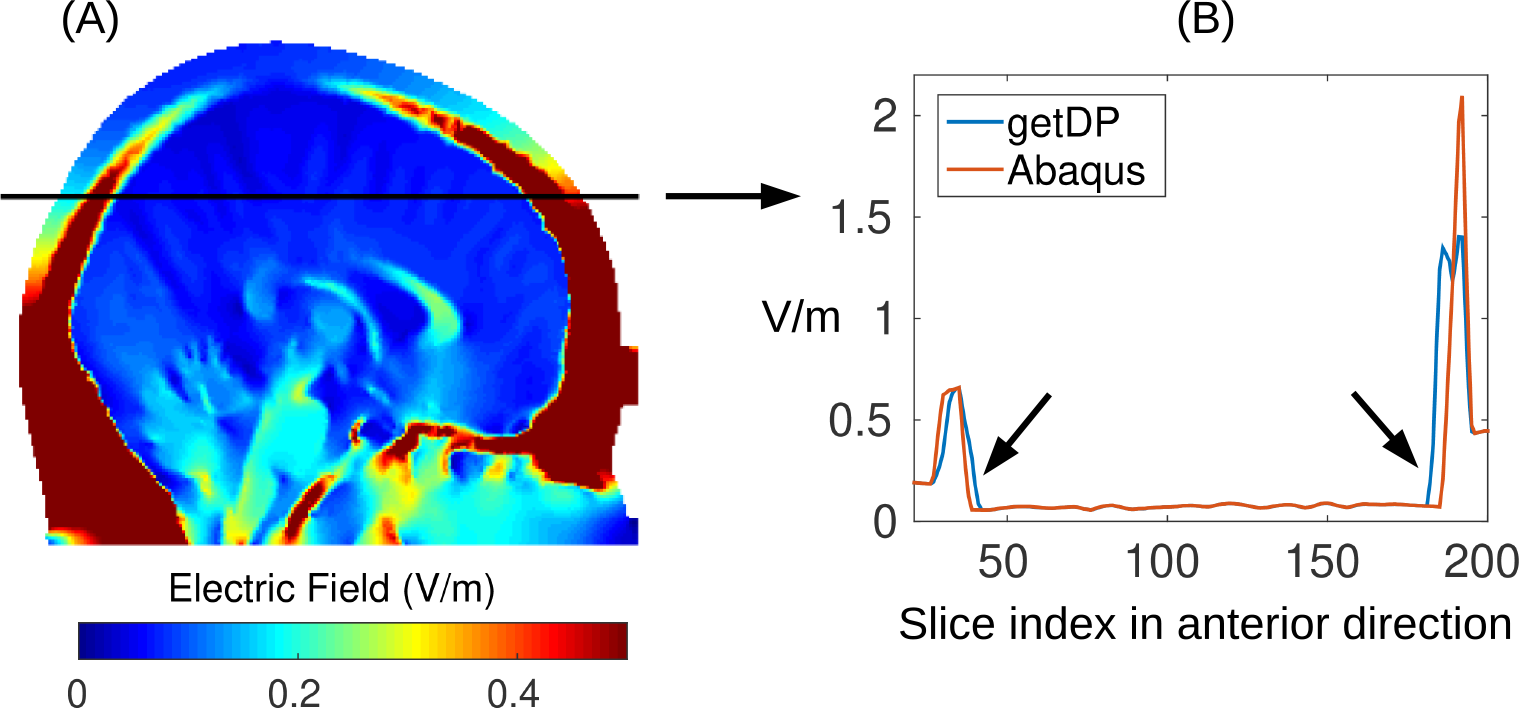
(A) Electric field magnitude in an example slice; (B) Field magnitude along the black line in (A) plotted for two different solvers: open-source getDP and commercial Abaqus. The results are based on an identical mesh, i.e., from using ROAST-sg and ROAST-sa in Figure 1. Note the electric field magnitudes are identical from two solvers except on the boundaries between brain, CSF and skull (indicated by the arrows in Panel B).

Comparisons between ROAST and SimNIBS on a larger dataset (Huang et al., 2016) is shown in Figure 6C. See Section 3.3 for details.

**Figure 6:**
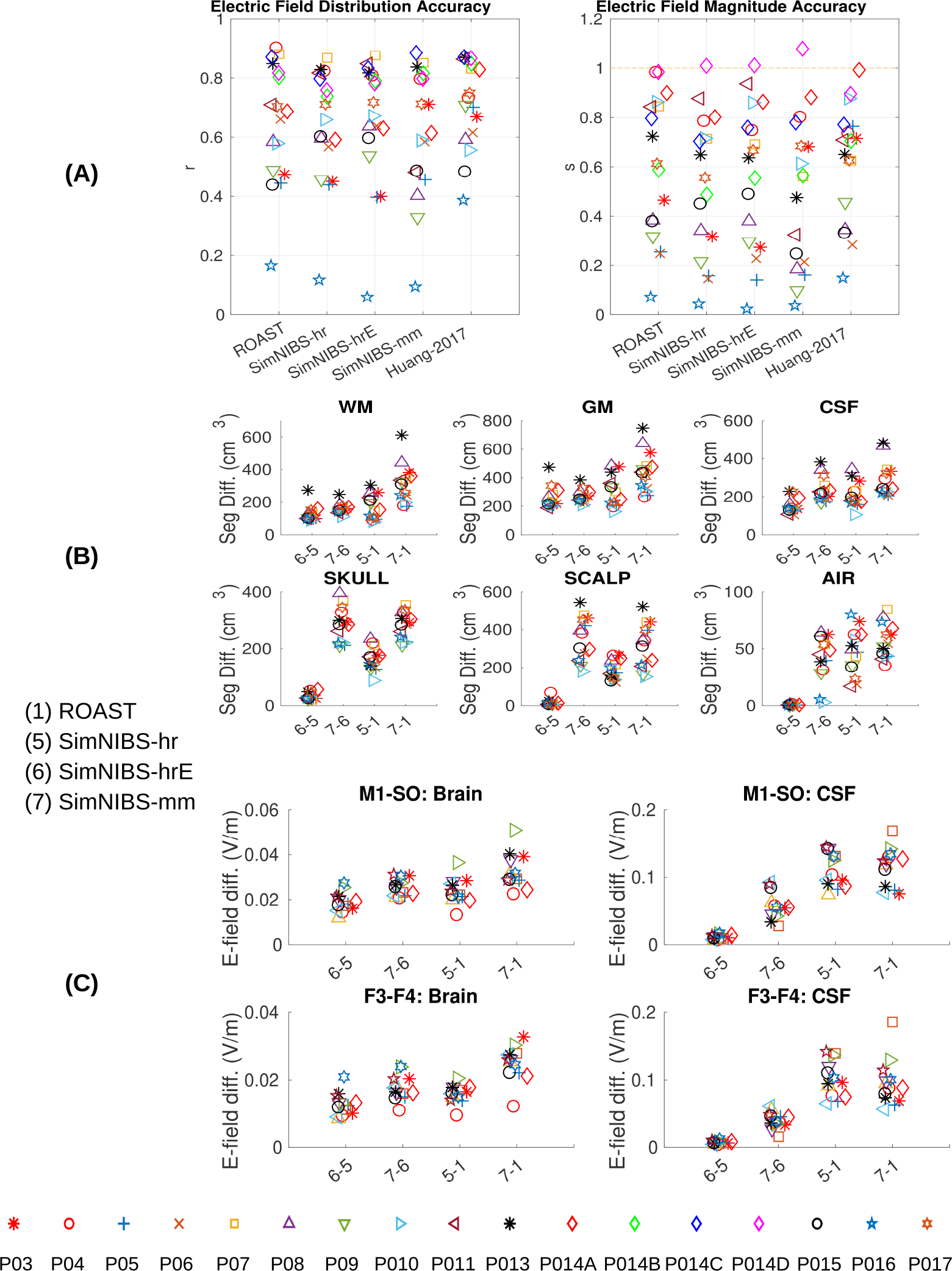
(A) Comparing the four modeling pipelines and an old published manual approach (Huang et al., 2017) using intracranial electrical recordings from human subjects under transcranial stimulation as the ground truth. Left panel in (A): Correlation indicates the accuracy of the spatial distribution. Right panel in (A): Scaling needed to best match the estimated fields with the measured fields. Correct magnitude prediction corresponds to a scale *s* = 1. Also the differences between the four pipelines in terms of segmentaion and electric field distributions are shown in (B) and (C).

### 3.3. Prediction accuracy of different modeling pipelines

The difference in the predicted electric fields begs the questions as to which of these pipelines best match observed electric fields recorded in humans. To address this we compared the predictions to intracranial field measurements recorded *in vivo* from 14 human subjects using publicly available data (Huang et al., 2016). In total we have 17 configurations (subject P014 has 4 stimulation montages) and a total of 1456 brain locations to compare against the models. In these recordings stimulation electrodes were placed on frontal and occipital poles to ensure maximal distance from the temporal craniotomies (Huang et al., 2017).

Figure 6A shows the results of four modeling pipelines – one for ROAST, three for SimNIBS. We also report results for models that we previously built by hand for these patients (Huang et al., 2017). These are included here to give a sense of the importance of implementing surgical details of these patients’ anatomies.

Performance is evaluated in terms of the correlation across recording locations between measurement and prediction (Figure 6A left panel) following Huang et al. (2017). This captures the accuracy of the predicted field distribution across the brain. Separate from that, we evaluate the accuracy of the field magnitude, as the scaling factor required for the predictions to best match the measured field magnitudes (Figure 6A right panel; scale of 1 corresponds to accurate estimation of magnitude, all models appear to over-estimate the magnitude).

There is no significant difference between ROAST and SimNIBS in predicting the electric field distribution (Figure 6A left panel, pairwise t-test: ROAST vs. SimNIBS-hr: *p* = 0.44; ROAST vs. SimNIBS-hrE: *p* = 0.95; ROAST vs. SimNIBS-mm: *p* = 0.17). However, ROAST predicts the electric field magnitude significantly better than SimNIBS-hr (Figure 6A right panel: *p* = 0.0003, *t*(16) = 4.60) and SimNIBS-mm (*p* = 0.009, *t*(16) = 2.97), and with a lesser difference from SimNIBS-hrE (*p* = 0.08). This means that the estimated slopes are somewhat closer to 1, although clearly all models over-estimate field magnitudes, which can be readily corrected if conductivities are calibrated to match the data (Huang et al., 2017). As expected, the manual approach outperforms the fully automated pipelines, including ROAST in terms of predicting the field distribution (Huang-2017 vs ROAST: *p* = 0.03, *t*(16) = 2.33). However, ROAST is no worse than the manually refined model (*p* = 0.62) in terms of estimating field magnitude.

As an additional point of reference, we also report the differences in segmentation (Figure 6B) and field estimates (Figure 6C) for all 14 subjects. Consistent with Section 3.1, we find that the different methods give similar segmentation results except for the contrast between FSL/FreeSurfer and SPM (7 vs. 6, 7 vs. 1). In terms of field intensities, the differences between these toolboxes in the brain are generally small, except for the old version of SimNIBS vs. the latest version of ROAST (7 vs. 1). The big difference in Figure 4D between SimNIBS-hr and ROAST-gg (5 vs. 4) drops for ROAST (5 vs. 1), indicating that in the lower grade of clinical MRI scans the differences between methods are less obvious.

## 4. Discussion and Conclusions

This paper reports on a new pipeline for current-flow modeling in the human head, which we have termed ROAST. Full-automation allows users to obtain state-of-the-art models based on individual MRIs without the know-how required to operate various complex software packages. The pipeline is based entirely on free software, except for Matlab, which we have retained in order to leverage existing tools (SPM, post-processing, electrode placement and 3D visualization). Installation is straightforward as all dependent libraries are included in a single package (for Linux, Windows, and Mac). ROAST allows for realistic modeling of the human head anatomy with relatively short run times (20–30 minutes) by leveraging efficient volumetric segmentation routines of SPM12. In the future it would be desirable to have a fully open-source solution for ROAST. The primary hurdle for this is the segmentation with SPM12 that runs on Matlab, which we may replace in future versions. More advanced user options to control segmentation may also be added. The current version of ROAST does not have a graphic user interface, and the visualization capability is limited compared to other software, especially SciRun. Future versions may improve visualization and user interface so that users can inspect the segmentation easily and perform any touch-up if needed. At present we recommend using free tools such as ITK-SNAP for this purpose (http://itksnap.org).

When comparing segmentation results to surface-based segmentation as implemented in FreeSurfer/FSL (SimNIBS-mm), it is clear that cortex is better segmented with surfaces capturing cortical foldings (Figure 2A), albeit at a substantial additional computational cost. On the other hand, volumetric segmentation implemented in SPM is relatively fast and has more flexibility in representing arbitrary morphology (Figure 2B–D under ROAST and SimNIBS-hr vol). Specifically, surface-based segmentation is limited to tissue volumes defined as the space between two surfaces (e.g., skull volume is between scalp and skull surface). Therefore, embedded structures (e.g., ventricles inside surface CSF), components with intersecting surfaces (e.g., gray matter and cerebellum), and tissues with disconnected regions (e.g., skull and disjoint spine vertebrae) cannot be defined with closed, non-intersecting surfaces, unless each structure is defined as separate surfaces. Note that we evaluated the segmentation only on one head because only this head has manual segmentation that was fully labeled by hand and we do not want to introduce any bias by using manual segmentation initialized by any automated segmentation routine. There are public MRI datasets (e.g. the Internet Brain Segmentation Repository (IBSR), LPBA40 (Shattuck et al., 2008), segmentation validation engine (Shattuck et al., 2009)) but none of them have manual segmentation of both brain and non-brain tissues which are needed for TES modeling.

When comparing the predicted fields between different toolboxes, one surprising finding is that the same volumetric segmentation approaches provide quite different results (Figure 4D, 5 vs. 4). One explanation is that SimNIBS converts the volumetric segmentation of SMP12 into surfaces prior to meshing, and in that process, many details of the anatomy are lost (Figure 2CD compare ROAST with SimNIBS-hr surf and SimNIBS-mm for the facial structures, optic canals, and the foramen magnum; also Figure 3). ROAST circumvents these problems by working entirely with volumes. However, in practice one does not seem to see this big difference (Figure 6C, 5 vs. 1, and Figure 6A). An additional question we wanted to address is whether replacing commercial software packages with free software tools affects the simulation results (iso2mesh and getDP instead of ScanIP and Abaqus). The difference turns out to be minor in the brain (~0.01 V/m, Figure 4D, Brain, 1 vs. 3). However, the difference are bigger in CSF (~0.1 V/m, Figure 4D, CSF, 1 vs. 3). This however is not unexpected given that these tools differ on how they compute electric fields from the potential distributions at the boundaries (Figure 5), where tissue conductivity values are discontinuous (Engwer et al., 2017). Despite nearly identical potential distributions the two methods give different field estimates at the boundaries. So in total, we conclude that the two methods give identical results, at least in terms of voltage distribution.

To validate predicted fields we compare them to field measured intracranially in human (Huang et al., 2016). For all methods the correspondence is moderate with correlations between measured and predicted fields in the range of 0.4 to 0.9 (Figure 6 A), except for one outlier patient (P016), where the detailed modeling of the surgical alterations may have been important. One surprising result of this analysis is that the performance of SimNIBS-mm is no worse than ROAST despite a severely limited FOV in the segmentation (missing neck and head below the nose). We interpret this finding as support for the notion that careful modeling of cortical gyrations, in particular CSF-filled sulci, is crucial to correctly predict field distribution in the brain. Neither ROAST nor SimNIBS are designed to capture the surgical details necessary to obtain accurate estimates on these intracranial field measurements. In these experiments where intracranial electric potentials were measured directly, the stimulating electrodes were placed far from the craniotomy. This was motivated by previous modeling studies that suggested minimal effects of skull defects on current flow for distant stimulation electrodes (Datta et al., 2010). Thus the hope was that the automated pipelines perform reasonably well on these data, despite lacking surgical details. In fact, in terms of field magnitude, ROAST is no worse than the manual models on this small sample. Overall, however, the caveat remains that this may not be the best dataset to compare different modeling approaches designed for intact anatomies. An alternative approach for validation might be to compare with recordings using stereotactic EEG. This recording method needs only minor burr-holes through the skull and allows more flexible placement of TES and EEG electrodes (Koessler et al., 2017). To validate models on intact anatomy one can also compare magnetic fields generated by TES to those measured with MRI (Jog et al., 2016; Göksu et al., 2018). However, additional work is needed in this field to improve signal quality.

Many of the recording electrodes in this study were placed on cortex (electrocorticogram electrodes) where field predictions are complicated because of the boundary issues discussed above. Since these recordings would be crucial to tease apart effects from surface vs. volume-based FEM modeling, the present results in Figure 6A should be explained with caution. New methods that promise to cope with conductivity discontinuities at boundaries (Engwer et al., 2017) should be tested and potentially incorporated into modeling pipelines. This may be particularly important for fields measured at the cortical surface.

The most notable finding of the experimental comparison in the present study is that for a number of subjects both manual and automated methods perform fairly well, and for other subjects, both seem to perform poorly. This poor performance may indicate inaccurate specification of the recordings setup, such as the locations of the stimulating electrodes. There may also be genuine inter-individual differences beyond what is captured by MRI-based segmentation, for example, tissue conductivity values may vary substantially between subjects. A detailed analysis of how modeling choices affect model performance on this data was presented in (Huang et al., 2017). Overall we feel that improving prediction accuracy will be an iterative process of improving experimental recording as well as modeling approaches.

Note that detailed knowledge of SimNIBS are needed for modeling some of the subjects in Figure 6A, also for one subject we had to exclude one outlier data point solved from SimNIBS, and one subject had to be excluded in Figure 6C as we could not fix all the issues from SimNIBS. On the other hand, ROAST is able to generate models for all these subjects out-of-box without any specific know-how for users.

Future theoretical work on novel modeling approaches is needed to advance this field further. For example, an FEM solver that can directly operate on segmentation data without first building a mesh (Nüßing et al., 2016). Additionally, more validation is needed for the FEM meshers and solvers using analytic solutions from a spherical model as the ground truth (Dmochowski et al., 2012), for which we provided a software interface (https://www.parralab.org/spheres/).

In closing we want to note the benefits of having multiple modeling toolboxes in development. For instance, SimNIBS has adopted SPM12 as the segmentation tool to allow faster volumetric segmentation. ROAST has in turn borrowed from SimNIBS the use of getDP as a free and fast FEM solver. Simalarly, it has been useful during the development of ROAST to have commercial-grade meshing and FEM solving tools as a reference point for our work (SimpleWare, Abaqus, COMSOL). We hope that future versions of these tools will continue benefiting from one another to maintain a healthy and productive software development community.

## ACKNOWLEDGMENT

The authors would like to thank Chris Thomas at Soterix for insightful discussions, and his work in marking up the special regions in the skull prior map for the patching algorithm in ROAST. We also would like to thank Alexander Opitz for pointing us to the free solver getDP. We also thank Jens Madsen for assistance with Matlab GUI programming, and Zeinab Esmaeilpour for testing ROAST extensively on the Mac OS. This work was supported by the NIH through grants R01MH111896, R44NS092144, R41NS076123, and by Soterix Medical Inc.

